# The Spike protein of SARS-CoV-2 impairs lipid metabolism and increases susceptibility to lipotoxicity: implication for a role of Nrf2

**DOI:** 10.1101/2022.04.19.488806

**Authors:** Vi Nguyen, Yuping Zhang, Chao Gao, Xiaoling Cao, Yan Tian, Wayne Carver, Hippokratis Kiaris, Taixing Cui, Wenbin Tan

**Affiliations:** Department of Cell Biology and Anatomy, School of Medicine, University of South Carolina, Columbia, South Carolina, USA; Department of General Surgery, The 3rd Xiangya Hospital of Central South University, Changsha, Hunan, China; Department of Obstetrics and Gynecology, Xiangya Hospital, Central South University, Changsha, Hunan, China; Department of Biomedical Engineering, College of Engineering and Computing, University of South Carolina, Columbia, South Carolina, USA; Drug Discovery & Biomedical Sciences, College of Pharmacy, University of South Carolina, Columbia, South Carolina, USA

**Keywords:** Spike protein, SARS-CoV-2, Lipid metabolism, Lipotoxicity, Autophagy, Ferroptosis

## Abstract

**Background/objectives:** Coronavirus disease 2019 (COVID-19) patients exhibit lipid metabolic alterations, but the mechanism remains unknown. In this study, we aimed to investigate whether the Spike protein of severe acute respiratory syndrome coronavirus 2 (SARS-CoV-2) impairs lipid metabolism in host cells.

**Methods:** A Spike cell line in HEK293 was generated using the pcDNA vector carrying the Spike gene expression cassette. A control cell line was generated using the empty pcDNA vector. Gene expression profiles related to lipid metabolic, autophagic, and ferroptotic pathways were investigated. Palmitic acid (PA)-overload was used to assess lipotoxicity-induced necrosis.

**Results:** As compared with controls, the Spike cells showed a significant increase in lipid depositions on cell membranes as well as dysregulation of expression of a panel of molecules involved lipid metabolism, autophagy, and ferroptosis. The Spike cells showed an upregulation of nuclear factor erythroid 2-related factor 2 (Nrf2), a multifunctional transcriptional factor, in response to PA. Furthermore, the Spike cells exhibited increased necrosis in response to PA-induced lipotoxicity compared to control cells in a time- and dose-dependent manner *via* ferroptosis, which could be attenuated by the Nrf2 inhibitor trigonelline.

**Conclusions:** The Spike protein impairs lipid metabolic and autophagic pathways in host cells, leading to increased susceptibility to lipotoxicity via ferroptosis which can be suppressed by a Nrf2 inhibitor. This data also suggests a central role of Nrf2 in Spike-induced lipid metabolic impairments.

**Highlights:**

- The Spike protein increases lipid deposition in host cell membranes
- The Spike protein impairs lipid metabolic and autophagic pathways
- The Spike protein exaggerates PA-induced lipotoxicity in host cells *via* ferroptosis
- Nrf2 inhibitor Trigonelline can mitigate the Spike protein-induced necrosis

## Introduction

Coronavirus disease 19 (COVID-19) is a pandemic viral infection that threatens global public health since the initial outbreak in December 2019 at the epicenter of Wuhan City, Hubei Province, China.^1^ COVID-19 is caused by the severe acute respiratory syndrome coronavirus 2 (SARS-CoV-2) with high pathogenicity and contagiousness.^2^ SARS-CoV-2 is a positive-sense single-stranded RNA virus that is capable of infecting human beings, together with six other coronaviruses.^2^ SARS-CoV-2 is assumed to be zoonotic and shares 96.3% sequence identity with the bat coronavirus RaTG13.^3^ The SARS-COV2 Spike protein is thought to be the receptor for the virus entry into host cells via surface angiotensin converting enzyme 2 (ACE2). Cleavage of Spike protein by Furin and Transmembrane Serine Protease 2 (TMPRSS2) facilitates SARS-CoV-2 entry into host cells^4^. ACE2 and TMPRSS2 are thought to be host determinants during initial infection.^5^ In addition, Neuropillin (NRP) 1 has been identified as the second host factor to facilitate SARS-CoV-2 entry upon cleavage by Furin.^6–8^

COVID-19 patients can be asymptomatic or symptomatic. Mortality rate of the COVID-19 varies in different geographic locations and patient populations.^1^ Patients with metabolic-associated preconditions such as hypertension, cardiovascular disorders (CVD), obesity and diabetes mellitus (DM) are susceptible to experience more severe symptoms.^9^ One common pathogenic co-factor related to hypertension, obesity, DM and CVD is hypercholesterolemia. Emerging evidence reveals direct interplay between COVID-19 and dire cardiovascular complications including myocardial injury and heart failure that are accompanied by elevated risk and adverse outcomes among infected patients.^10^ COVID-19-associated cardiac complications may become even worse in the setting of cardiometabolic pathologies associated with obesity, although obesity per se is a strong risk factor for severe COVID-19.^11^ Our recent studies have shown decreased levels of total cholesterol (TC), low density lipoprotein cholesterol (LDL-c) and high density lipoprotein cholesterol (HDL-c) in COVID-19 patients, which are associated with disease severity and mortality.^12–14^ Mechanistically, lipids have been shown to be a critical contributor to transmission, replication and transportation for some types of viruses. For example, lipid rafts have been reported to be necessary for SARS virus replication.^15^ Although it has been firmly established that obesity and obesity-related complications are major risk factors for COVID-19 severity, the underlying mechanisms have yet to be determined. In this study, we attempted to explore the direct interplay between the Spike protein and lipid metabolism pathways in host cells. We found that the Spike protein impairs lipid metabolic and autophagic pathways in host cells, leading to increased susceptibility to lipotoxicity most likely via switching on nuclear factor erythroid 2-related factor 2 (Nrf2)-mediated ferroptosis.

## Methods

### Materials

Human embryonic kidney 293 cells (HEK293) and DMEM medium were purchased from ATCC (Manassas, VA, USA). Coding sequence of SARS-CoV-2 S gene (GenBank: QHU36824.1) fusion with a c-terminal His tag was synthesized *in vitro* (Genscript, Piscataway, NJ, USA) after an optimization for expression in human cells.^16^ The sequence was cloned into a pcDNA3.1 vector to obtain pcDNA-Spike.^16^

Anti-His tag and microtubule-associated protein 1 light chain 3 beta (LC3) Antibodies were purchased from Millipore Sigma (Burlington, MA, USA). Anti-Adipose Differentiation-Related Protein (ADRP, or Perilipin-2, PLIN2), Nrf2, prostaglandin E synthase 2 (PTGS2), and PI3K-beta antibodies were obtained from ProteinTech (Rosemont, IL, USA). Anti-ATG7 antibody was purchased from Abcam (Waltham, MA, USA). Anti-SRB1 antibody was obtained from Novus (Centennial, CO, USA). Anti-Fth1, HRP-anti-rabbit or mouse secondary antibodies, and RIPA lysis buffer were obtained from Santa Cruz Biotech (Dallas, TX, USA). Primers for real time RT-PCR were synthesized by IDT (Coralville, IA, USA) (Supplementary table 1). RNA extraction kit was obtained from Zymo Research (Irvine, CA, USA). The RT kit was obtained from Takara Bio USA (Moutain view, CA, USA). The SYBR green master mix was from Biorad (Hercules, CA, USA).

### Generation of the Spike-protein stable expression cell line

The HEK293 cells were grown in DMEM containing 10% FBS and transfected by pcDNA-Spike or pcDNA vector using lipofectamine 300 reagent (ThermoFisher, Waltham, MA, USA). Two days after transfection, the cells were treated by G418 starting from the concentration of 100 μg/ml with a gradual increase to 800 μg/ml during the following 2 weeks. The individual colonies with stable integration of the pcDNA-Spike (HEK_Spike) or pcDNA vector (HEK_pcDNA) were selected and expanded. The HEK_Spike stable colonies were confirmed to express the Spike protein using immunoblot. The cells were maintained in DMEM with10% FBS regularly for further experiments.

### Lipid (Oil Red O) staining

The Lipid (Oil Red O) kit was obtained from Sigma-Aldrich (St. Louis, MO, USA). The HEK293, HEK_pcDNA, and HEK_Spike cells were fixed by 10% formalin and followed by 60% isopropanol treatment. The Oil Red O working solution was added into the cells and incubated for 15 minutes. The cell nucleus was counterstained using hematoxylin. The images were acquired using an ImagXpress Pico Automated system (Molecular Device, San Jose, CA, USA). For the quantification of Oil Red O staining, stain was extracted in isopropanol and absorbance was measured at 492 nm using a Thermo Scientific Multiskan Spectrophotometer system (ThermoFisher, Waltham, MA, USA).

### Real time-RT PCR and Western blot

To generate cDNA, 1.0~5 μg of total RNA was reverse-transcribed in a 20-μl reaction containing 1x RT buffer (Clontech, Mountain View, CA, USA), 0.5 mM dNTPs, 0.5 μg of oligo (dT) 15-mer primer, 20 units of RNasin, and 5 units of SMART Moloney murine leukemia virus reverse transcriptase (Takara Bio, Mountain View, CA, USA). The RT reaction was carried out at 42°C for 2 hrs. A converted index for three house-keeping genes were used to normalize the amplification data: GAPDH, Nono, and β-actin. Expression levels of a panel of 83 genes related to lipid metabolic, autophagic, and ferroptotic pathways (Supplementary Table 1) were determined using real time RT-PCR. The reaction for the multiplex real time PCRs contained 1× SYBR Green qPCR Master Mix (Bio-Rad, Hercules, CA, USA), 10 ng of each template, and 10 pmol of each specific primer in a 25-μl total volume in a 96-well format. Each reaction was performed in duplicate under identical conditions. The PCR conditions were one cycle at 95°C for 2 min followed by 45 cycles of 15 s at 95°C and 60 s at 60°C. Relative quantification of the real time PCR was based upon the amplification efficiency of the target and reference genes and the cycle number at which fluorescence crossed a prescribed background level, cycle threshold (*Ct*).

For immunoblot assay, cell lysates were extracted from HEK, HEK_pcDNA, or HEK_Spike cells using RIPA lysis buffer (Santa Cruz Biotech., Inc., Dallas, TX, USA). Proteins were separated by SDS-PAGE and transferred onto PVDF membranes. Primary antibodies were used to detect the expression of specific proteins and followed by HRP-labeled secondary antibodies. Images were acquired using a Bio-Rad Gel Imaging System (Hercules, CA, USA).

### Palmitic acid (PA)-induced lipotoxicity assay

The HEK293, HEK_pcDNA and HEK_Spike cells were cultured in a 96-well plate and reached 80% confluence on the next day before treatment. The cells were kept in DMEM medium with 5% FBS during the entire treatment procedure. The cells were treated with PA in various concentrations from 250 to 1000 μM for 24, 48, or 72 hours. For treatment control groups, the cells were treated by the same concentrations of BSA in parallel. For the inhibitory assay, the cells were pre-incubated by autophagic inhibitors (125 μM trigonelline (TRG), or 10 μM Wortmannin), or a ferropotosis inhibitor (ferrostatin, 2 μM), or an apoptosis inhibitor (necrostatin-1, 10 μM) in DMEM medium with 5% FBS for 2 hours, followed by 750 μM PA treatment for 24 hours. The cells were then co-stained by Propidium Iodide (PI) and Hoechst 33342 NucBlue Live Cell Stain dye (ThermoFisher, Waltham, MA, USA) to show the dead and live cells. The dyes were excited at 535 nm and 405 nm, respectively. Images were acquired and cell viability analyses were performed using an ImagXpress Pico Automated system (Molecular Device, San Jose, CA, USA).

### Viral production and H9C2 cell culture

The Spike gene was cleaved from pcDNA-Spike plasmid and cloned into lentiviral vector pLV-mCherry (Addgene, Watertown, MA, USA) with removal of mCherry gene to generate pLV-Spike plasmid. The lentiviral production followed our previous report with slight modifications.^16^ We generated a Spike-pseudotyped (Spp) lentivirus with the Spike protein as the viral surface tropism as well as expression of the Spike protein, which was referred to as Cov-Spp-S virus. Briefly, Phoenix cells (ATCC, Manassas, VA, USA) were cultured in a Dulbecco’s Modified Eagle’s medium (DMEM) containing 10% FBS and transfected by pLV-Spike using a calcium phosphate kit (ThermoFisher, Waltham, MA, USA). The control virus with VSV-G as the tropism and expression of mCherry was generated by co-transfection of pLV-mCherry and pMD2.G vector (Addgene, Watertown, MA, USA) into the Phoenix cells, which was referred to as VSV-G virus. The supernatant with produced virus (Cov-Spp-S or VSV-G lentivirus) was harvested 72-hours post transfection, clarified by centrifuging at 5000 g for 15 min followed by filtration of the supernatant through a 0.45 μm filter disk. The virus was collected by ultracentrifugation at 24,000 rpm for 2 hours using Beckman SW41 rotor. The viral pellets were resuspended by cold PBS buffer and stored at −80 °C before use. The viral particle number was determined using a real time RT-PCR assay to quantify the RNA copies of the Spike or mCherry.

H9C2 cells (ATCC, Manassas, VA, USA) were cultured in DMEM (10% FBS) medium in a 96 well-plate with 80% confluence. The cells were infected twice at the first and second days using Cov-Spp-S or VSV-G lentivirus (4.8×10^7^ particles per well) with addition of polybrene (1 μg/ml); the cells were then incubated for an additional 5 days, followed by treatment with PA (675 μM) for 24 hours in DMEM medium containing 5% FBS. The same concentration of BSA was used for treatment control groups in parallel. For the inhibitory assay, the cells were pre-incubated with 125 μM TRG in DMEM medium with 5% FBS for 2 hours, followed by 675 μM PA treatment for 24 hours. The cells were then co-stained with Propidium Iodide (PI) and Hoechst 33342 NucBlue Live Cell Stain dye (ThermoFisher, Waltham, MA, USA) to show the dead and live cells, respectively.

### Statistical analyses

All statistical analysis were performed in Origin 2019. The student *t* test or one-way ANOVA was used for two groups or multiple comparisons test. The data was presented as “mean ± s.d.” and *p* < 0.05 was considered as significant.

## Results

The stable cell line with expression of the Spike protein was obtained upon neomycin selection after transfection of pcDNA_Spike into HEK293 cells. The control stable cell line with integration of the empty pcDNA was generated simultaneously. The expression of the Spike protein was verified using an anti-His tag antibody (Fig 1A). The same pcDNA_Spike vector was used for production of the Spike protein-pseudotyped (Spp) lentivirus in HEK Phoenix cells in our previous study where the expression of the Spike protein was verified using both anti-Spike S1 and S2 subunit antibodies.^16^ We did not observe a significant difference in growth rate between pcDNA and Spike stable cell lines (data not shown). To explore whether the Spike protein had a role in lipid metabolisms in host cells, we performed Oil Red O staining among these cell lines. We found there was a significant accumulation of lipid deposition in the Spike cells compared to pcDNA and mock control cells (Fig 1B). The histological staining showed that the lipid depositions were mainly located on the cell membrane in the Spike cell line (Fig 1C-E), indicating an impairment of lipid metabolism in host cells.

**Fig 1.**
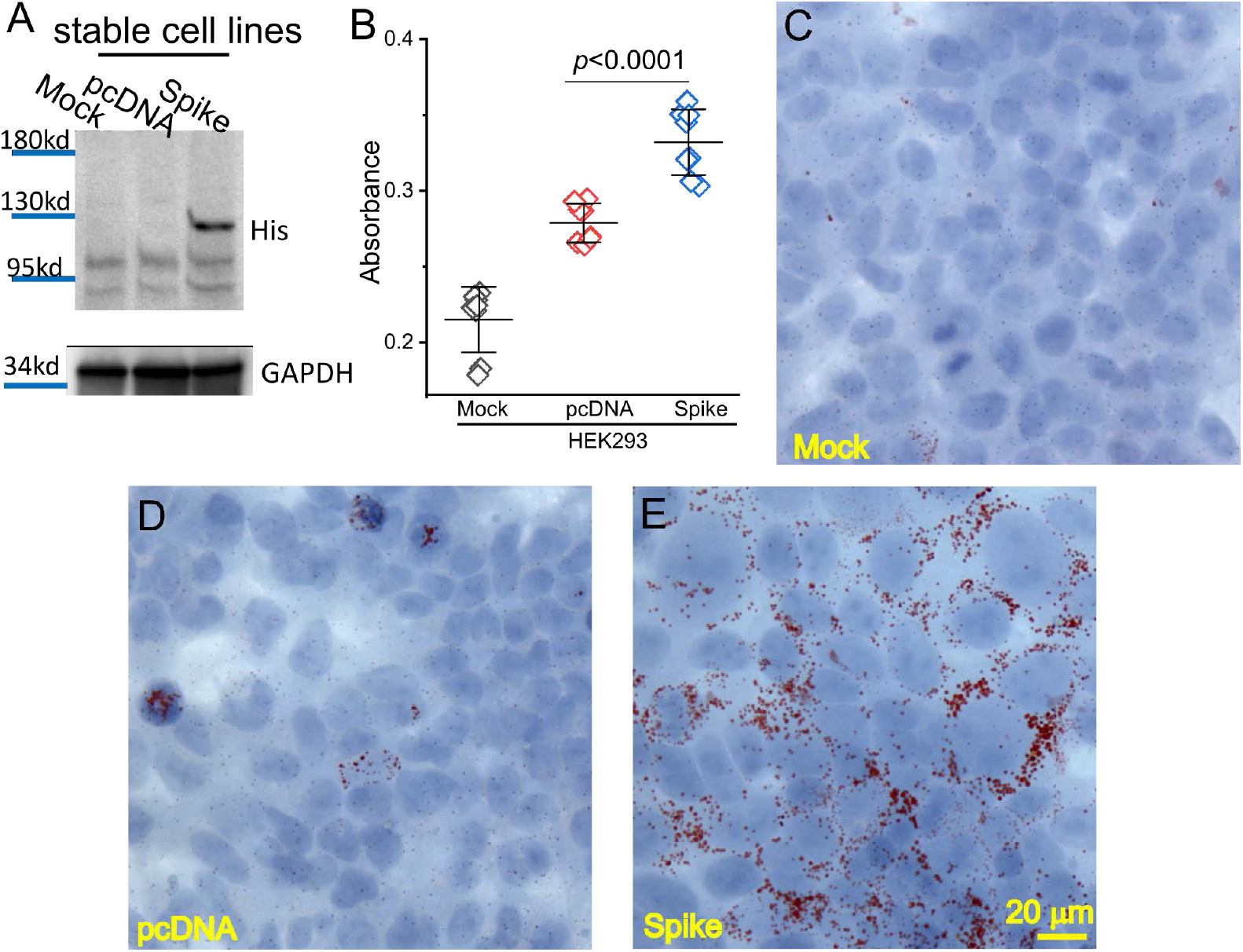
Spike protein caused lipid deposition in host cells. (A) Generation of Spike protein stable cell line. The expression of Spike protein was verified using an anti-His tag antibody in the Spike cells. (B) The quantification of Oil Red O staining in the mock, pcDNA, and Spike cells with measurement of an absorbance at 492 nm. (C-E) Histological images showing Oil Red O staining of lipid droplets in the mock (C), pcDNA (D), and Spike (E) cells. Nuclear components were counterstained by hematoxylin.

We next examined transcriptional levels of a panel of 83 genes that are representative markers of lipid metabolism, autophagy, and ferroptosis (Supplementary Table 1). We found the mRNA levels of many lipid metabolic markers such as proprotein convertase subtilisin/kexin type 9 (Psck9), SREBF chaperone (Scap), Plin2, low density lipoprotein receptor-related protein 10 (Irp10), lecithin-cholesterol acyltransferase (lcat), low density lipoprotein receptor-related protein associated protein 1 (Irpap1), oxysterol binding protein-like 5 (Osbpl5), oxysterol binding protein-like 1A (Osbpl1a), protein kinase, AMP-activated, gamma 2 non-catalytic subunit (Prkag2), and mevalonate kinase (Mvk) were upregulated while Serpinb2 was downregulated in the Spike cells (Fig 2A). For the examined autophagic and ferroptotic markers, Atg3, Atg7, Atg12, Nrf2, Phosphatidylinositol-4,5-Bisphosphate 3-Kinase Catalytic Subunit Alpha (Pik3ca), Pik3cd, Phosphoinositide-3-Kinase Regulatory Subunit 3 (Pik3r3), acyl-CoA synthetase long-chain family 4 (Acsl4), ferritin heavy chain (Fth1), glutaminase 2 (Gls2), and Ptgs2 showed an increase in mRNA levels in the Spike cells; while Pik3c3, Atg5, and Hamp showed a decrease in mRNA levels in the Spike cells (Fig 2B, C). These results reveal a negative impact of Spike proteins per se on lipid metabolism, autophagy, and probably ferroptosis in the cell.

**Fig 2.**
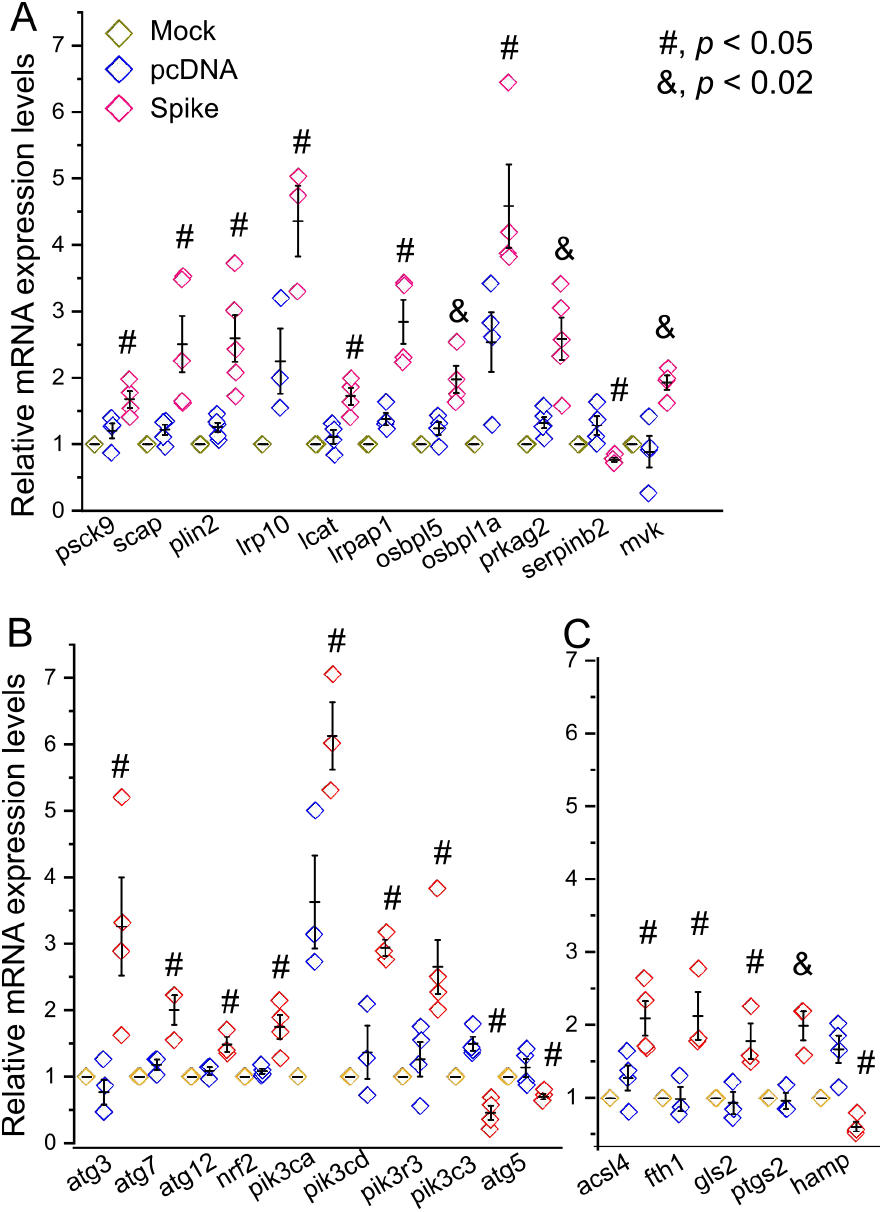
Transcriptional profiles of representative biomarkers for lipid metabolism, autophagy, and ferroptosis. Relative mRNA levels of biomarkers involving lipid metabolism (A), autophagy (B), and ferroptosis (C) in the mock, pcDNA, and Spike cells. The normalization was performed using a converted index of β-actin, nono, and GAPDH as the reference genes.

We further examined the biological significance of Spike protein expression with a focus on lipotoxicity in vitro. PA overload induced cell death in cultured HNK293 and pcDNA cells (Fig 3, Supplementary Fig 1), demonstrating a PA-induced lipotoxicity as described elsewhere.^17^ However, the PA-induced cell death was significantly augmented in Spike cells (Fig. 3, Supplementary Fig. 1). The PA-induced cell death also showed a dose- and time-dependent response (Fig 3, Supplementary Fig 1). The BSA control did not cause any significant cell death among these cell lines (Supplementary Fig 2). We next examined the expression patterns of some crucial factors in response to PA-induced lipotoxicity. SRB1, ATG7, PTGS2, showed a significantly higher level in the Spike cells than the mock and pcDNA control cells (Fig 4), which were consistent with the changes in mRNA levels (Fig 2). In response to PA treatment, the levels of SRB1, ATG7, PTGS2, LC3 I/II ratio, Fth1, in Spike cells increased significantly as compared with the pcDNA control cells (Fig 4).

**Fig 3.**
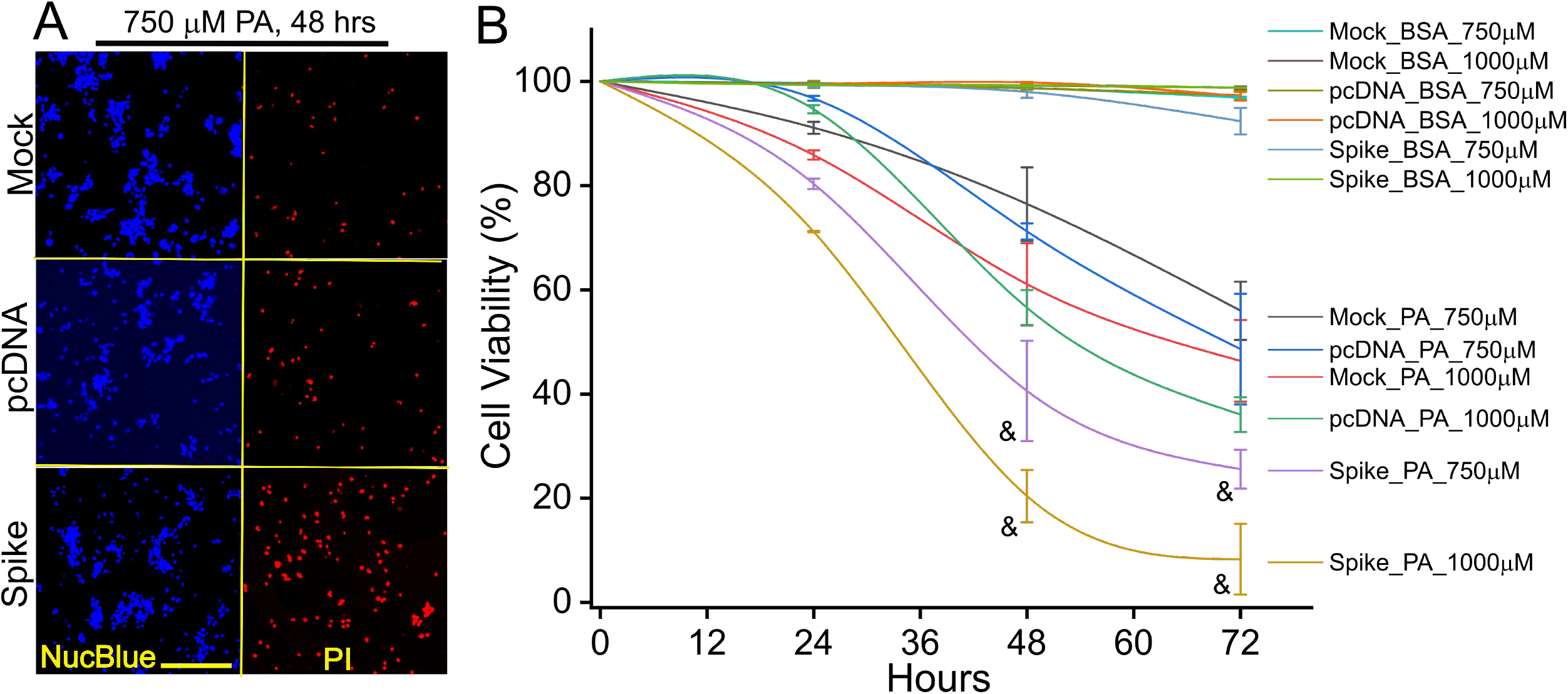
The Spike protein-exaggerated PA-induced lipotoxicity in host cells. (A) Fluorescent images showing the cell viability in the mock, pcDNA, and Spike cells treated by 750 μM PA for 48 hours in DMEM containing 5% FBS. The cells were co-stained by Propidium Iodide (PI) and Hoechst 33342 NucBlue Live Cell Stain dye to show the dead and live cells, respectively. Scale bar, 25 μm. (B) Cell viability curves of mock, pcDNA, and Spike cells in response to PA (750 or 1000 μM) treatments for 24 to 72 h. &, indicates *p*<0.05 in the Spike cells as compared with the mock and pcDNA control cells in response to the same dosages of PA at 48 or 72 h. n=5~6 wells in each dose and time point.

**Fig 4.**
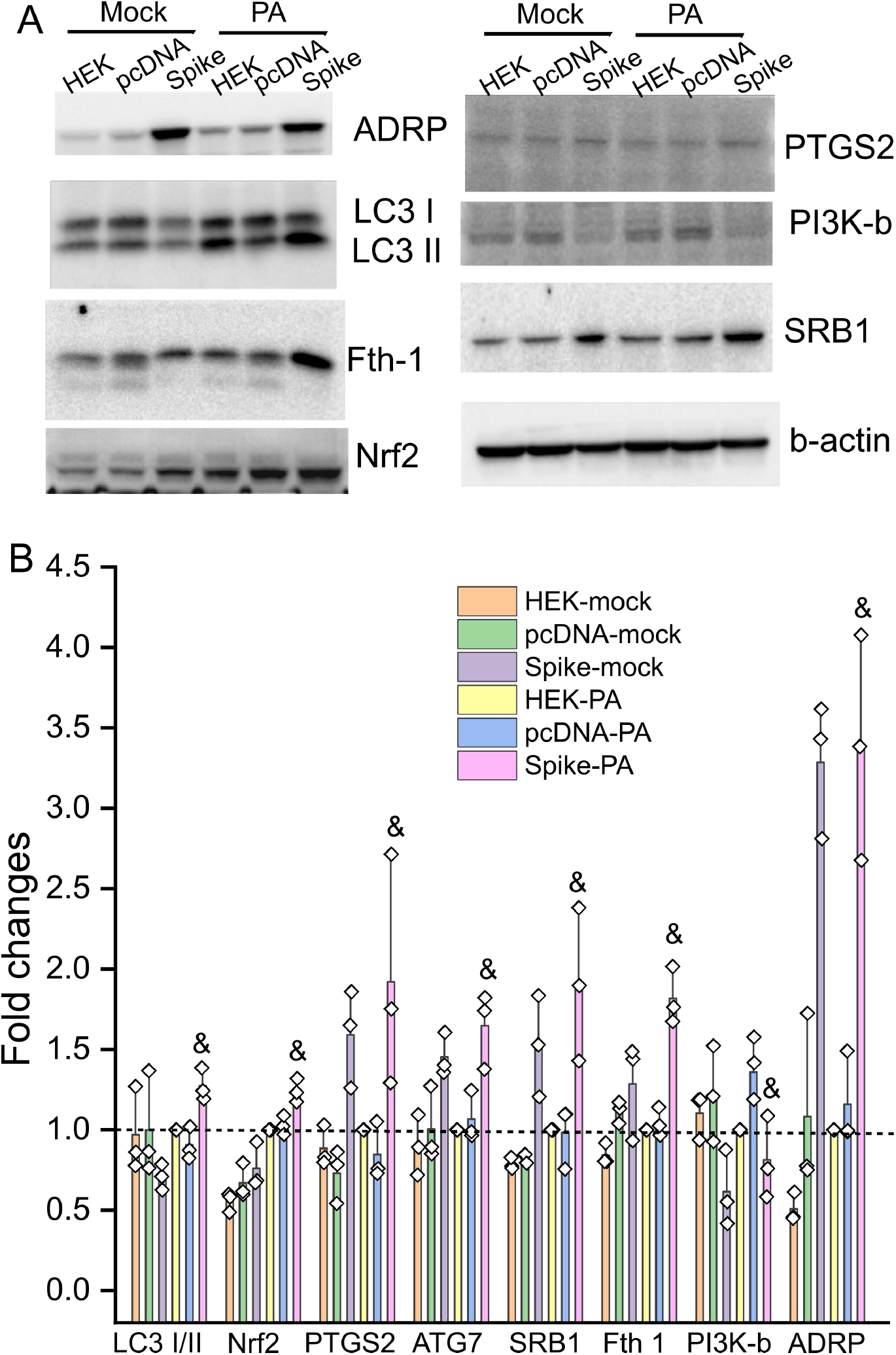
Immunoblot showing the relative changes in levels of some representative biomarkers of impaired lipid metabolism, autophagy, and ferroptosis in response to PA overload. The mock, pcDNA, and Spike cells were treated by 750 μM PA for 24 hours in DMEM containing 5% FBS. (A) Representative immunoblot showing the protein levels of ADRP, LCI/II, Fth1, Nrf2, PTGS2, PI3K-b, SRB1, and beta-actin in the mock, pcDNA, and Spike cells with or without PA treatment. (B) The relative quantification of each biomarker in cells. The protein level for each biomarker was first normalized to the corresponding beta-actin level. The relative ratio of each biomarker within group was then obtained using its’ level in the mock group with PA treatment as 1 (indicated by the dashed line). &, indicates *p*<0.05 in the Spike cells as compared with the pcDNA control cells with the PA treatment. n=3 independent experiments.

We have previously shown that Nrf2 is the crucial transcriptional factor mediating PA-induced ferroptosis in autophagy-impaired in cardiomyocytes under obese conditions.^18^ PI3K has been shown to play a critical role in autophagy.^19–21^ We then asked whether inhibitors for Nrf2, PI3K, and ferroptosis could attenuate PA-induced lipotoxicity in the Spike cells. Indeed, the Nrf2 inhibitor TRG, PI3K pan inhibitor Wortmannin, and ferroptosis inhibitor ferrostatin, but not a necroptosis inhibitor necrostatin 1, significantly mitigated the PA-induced Spike protein-exaggerated lipotoxicity (Fig 5).

**Fig 5.**
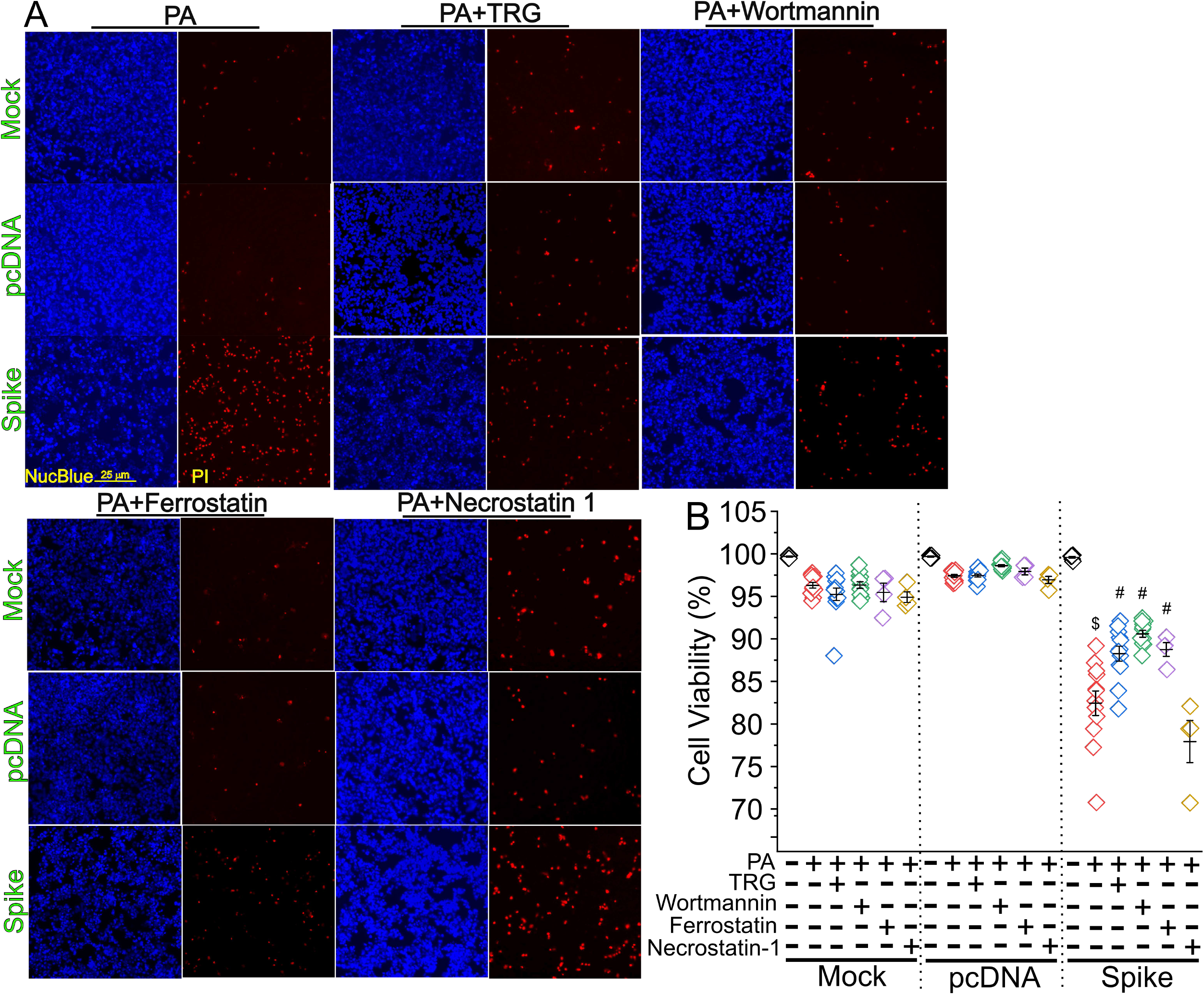
Autophagic/ferroptotic inhibitors attenuated the Spike protein-exaggerated PA-induced necrosis. (A) Fluorescent images showing the cell viability in the mock, pcDNA, and Spike cells treated by 750 μM PA treatment for 24 hours in DMEM containing 5% FBS with or without a Nrf2 inhibitor TRG, pan-PI3K inhibitor Wortmannin, ferroptosis inhibitor Ferrostain, and apoptotic inhibitor Necrostatin 1. The cells were co-stained by Propidium Iodide (PI) and Hoechst 33342 NucBlue Live Cell Stain dye to show the dead and live cells, respectively. (B) Quantitative data showing the cell viability in each treatment group. $, indicates *p*<0.05 in the Spike cells treated by PA as compared with the Spike cells without PA treatment. #, indicates *p*<0.05 in each group treated with PA and various inhibitors as compared with the Spike cells treated with PA only.

Finally, we attempted to test whether a similar mechanism could be recapitulated in H9C2 cells, a cardiomyocyte-like cell line. We infected the H9C2 cells using CoV-Spp-S lentivirus expressing the Spike protein or VSV-G control virus expressing mCherry. The cells infected by the CoV-Spp-S showed a significantly higher cell death than the cells being infected by the VSV-G control virus (Supplementary Fig 3); the PA-induced Spike protein-exaggerated lipotoxicity could be reversed using the Nrf2 inhibitor TRG (Supplementary Fig 3).

## Discussion

The data in this study has demonstrated that the Spike protein alone can directly impair lipid metabolic and autophagic pathways in host cells, leading to increased lipotoxicity through ferroptosis. This result has shown a direct and evident role of the Spike protein in exaggeration of pre-existing lipotoxicity, revealing a mechanistic insight into the clinical manifestations of high susceptibility and mortality rate of obese patients with COVID-19. Furthermore, we have shown that the Spike protein-induced necrosis can be suppressed by PI3K pan inhibitor Wortmannin, ferroptosis inhibitor ferrostatin 1, and Nrf2 inhibitor trigonelline. Trigonelline, an alkaloid enriched in coffee, is among those effective inhibitors, providing a potential and feasible preventive strategy to mitigate COVID-19-associated cardiometabolic pathologies associated with obesity.

Numerous studies have reported lipidomic dysregulation in COVID-19 patients. For example, Shen et al. showed a strong downregulation of over 100 lipids including sphigolipids, glycerophospholipids, fatty acids, and various apolipoproteins in COVID-19 patients.^22^ Increased levels of sphingomyelins (SMs), non-esterified fatty acids (NEFAs), and free polyunsaturated fatty acids (PUFAs) have been shown in COVID-19 patients as well.^23–25^ Increases in PLA2 activation, which results in long-chain PUFAs, may be associated with the COVID-19 disease deterioration.^26,27^ Our previous studies together with other reports have shown the downregulation of serum LDL-c and HDL-c levels in COVID-19 patients.^12–14,28^ These lines of accumulated evidence have revealed a central role of lipids and lipid metabolism in the development of COVID-19 disease. In this study, we demonstrate that the Spike protein executes a direct function in altering lipidome via upregulation of a panel of genes involving lipid metabolism and resulting in enhanced lipid deposition on the cell membrane. This data provides direct evidence showing that the Spike protein modulates lipid metabolism in host cells and is an important independent factor contributing to the altered lipidome in COVID-19 patients.

PI3Ks play important roles in autophagy formation during early stages of viral infection for both canonical and non-canonical endocytic pathways.^19–21^ They are also critical downstream components of growth factor receptor (GFR) signaling cascades which drive phosphorylation of viral proteins upon SARS-CoV-2 infection.^29^ Therefore, this class of enzymes has been proposed as a druggable target for prevention and treatment of SARS-CoV-2 infection. Indeed, inhibition of class I or class III PI3K prevents viral replication,^29,30^ probably through distinct mechanistic actions on different stages of SARS-CoV-2 viral life cycle. In a distinct mechanism, our data shows that the Spike protein alone can dysregulate expression of various PI3Ks in host cells including upregulation of class I PIK3CA, PIK3CD, and PIK3R3, but downregulation of class 3 PIK3C3 which is required for autophagosome and lysosome fusion^31^. In addition, pan-PI3K inhibitor wortmannin shows a potent inhibition of the Spike-protein exaggerated PA-induced lipotoxicity, suggesting that increasing autophagosome formation while decreasing autophagosome fusion with lysosomes thereby leading to accumulation of autophagosomes may be a cause of Spike protein-exaggerated lipotoxicity. Therefore, targeting of PI3K can be potentially beneficial for those COVID-19 patients with a metabolic precondition of hyperlipidemia.

The transcription factor Nrf2 controls the basal and induced expression of more than 1,000 genes in cells that can be clustered into several groups with distinct functions, such as antioxidative defense, detoxification, protein degradation, iron and lipid metabolism.^18^ Thus, the functions of Nrf2 spread rather broadly from antioxidative defense to protein quality control and metabolism regulation. Studies have demonstrated that Nrf2 is required for cardiac adaptation when cardiac autophagy is intact; however, it operates a pathological programme to exacerbate maladaptive cardiac remodeling and dysfunction when myocardial autophagy is inhibited in the settings of sustained pressure overload^32^ and chronic type 1 diabetes.^33^ Notably, chronic obesity, a pre-type 2 diabetic setting, results in inhibition of myocardial autophagy, thereby leading to cardiac pathological remodeling and dysfunction.^34,35^ In this study, the protein level of Nrf2 is upregulated in the Spike cells in response to PA treatment. Furthermore, TRG, a Nrf2 inhibitor, can attenuate Spike-protein-exaggerated PA-induced necrosis in the cell with impaired autophagy. Collectively, it is reasonable to posit a central role of Nrf2 in PA-induced Spike protein-exaggerated lipotoxicity in host cells. However, the detailed pathological mechanism and molecular interactions mediated by impaired Nrf2 pathways need further validation, which will be the goal of our future studies.

There are several questions yet to be answered, which will be the focus of our future studies. First, the types of altered lipids or lipid metabolites caused by the Spike protein have not been identified. Second, the molecular mechanism underlying the Spike protein-induced lipidomic dysregulation is not well understood. Our data indicate that Nrf2 may play a central role in the transcriptional level for this pathological process. However, this needs further elucidation and validation. Third, the current emerging variants, omicron strains, present multiple mutations in the Spike protein; whether these omicron versions of Spike protein variants can enhance or attenuate their functions in lipid metabolic alteration as compared with the alpha version of the Spike protein is unknown. Fourth, the Spike protein-induced impairments in both autophagic and lipid metabolic pathways in host cells are evident. However, whether autophagic impairment is a consequence of, or in parallel to lipid metabolic impairments is unknown. Most likely, autophagic impairment is intertangled with lipidomic alterations not only as a result but also an independent factor for the deterioration in response to lipotoxicity.

In conclusion, this study has demonstrated that the Spike protein can cause lipid deposition, impair lipid metabolic and autophagic pathways in host cells, ultimately leading to increased susceptibility to lipotoxicity via ferroptosis. The Spike protein-enhanced lipotoxicity can be suppressed by the Nrf2 inhibitor TRG, indicating a central role of Nrf2 in COVID-19-associated cardiac complications involving obesity.

## Acknowledgement

We greatly appreciate Dr. Mitzi Nagarkatti from the Department of Pathology, Microbiology and Immunology at University of South Carolina School of Medicine for their very kind and generous help with real time PCR assays. We are also very thankful to the support and assistance from Instrumentation Resource Facility at University of South Carolina School of Medicine. We are very grateful to the support from Dr. Igor Roninson and the COBRE Center for Targeted Therapeutics at University of South Carolina.

## References

1 CoronavirusResourceCenter. COVID-19 Dashboard by the Center for Systems Science and Engineering (CSSE) at Johns Hopkins University (JHU), <https://coronavirus.jhu.edu/map.html> (2020).

2 The species Severe acute respiratory syndrome-related coronavirus: classifying 2019-nCoV and naming it SARS-CoV-2. Nat Microbiol, doi:10.1038/s41564-020-0695-z (2020).

3 Zhou, P. et al. A pneumonia outbreak associated with a new coronavirus of probable bat origin. Nature, doi:10.1038/s41586-020-2012-7 (2020).

4 Walls, A. C. et al. Structure, Function, and Antigenicity of the SARS-CoV-2 Spike Glycoprotein. Cell 181, 281–292 e286, doi:10.1016/j.cell.2020.02.058 (2020).

5 Ziegler C et al. SARS-CoV-2 receptor ACE2 is an interferon-stimulated gene in human airway epithelial cells and is enriched in specific cell subsets across tissues. Cell, doi:DOI: 10.1016/j.cell.2020.04.035 (2020).

6 Daly, J. L. et al. Neuropilin-1 is a host factor for SARS-CoV-2 infection. Science, doi:10.1126/science.abd3072 (2020).

7 Cantuti-Castelvetri, L. et al. Neuropilin-1 facilitates SARS-CoV-2 cell entry and infectivity. Science, doi:10.1126/science.abd2985 (2020).

8 Moutal, A. et al. SARS-CoV-2 Spike protein co-opts VEGF-A/Neuropilin-1 receptor signaling to induce analgesia. Pain, doi:10.1097/j.pain.0000000000002097 (2020).

9 Richardson, S. et al. Presenting Characteristics, Comorbidities, and Outcomes Among 5700 Patients Hospitalized With COVID-19 in the New York City Area. Jama, doi:10.1001/jama.2020.6775 (2020).

10 Samidurai, A. & Das, A. Cardiovascular Complications Associated with COVID-19 and Potential Therapeutic~Strategies. Int J Mol Sci 21, doi:10.3390/ijms21186790 (2020).

11 Stefan, N., Birkenfeld, A. L. & Schulze, M. B. Global pandemics interconnected - obesity, impaired metabolic health and COVID-19. Nat Rev Endocrinol 17, 135–149, doi:10.1038/s41574-020-00462-1 (2021).

12 Fan, J. et al. Low-density lipoprotein is a potential predictor of poor prognosis in patients with coronavirus disease 2019. Metabolism, 154243, doi:10.1016/j.metabol.2020.154243 (2020).

13 Wei, X. et al. Hypolipidemia is associated with the severity of COVID-19. J Clin Lipidol, doi:doi: 10.1016/j.jacl.2020.04.008 (2020).

14 Cao, X., Yin, R., Albrecht, H., Fan, D. & Tan, W. Cholesterol: A new game player accelerating endothelial injuries caused by SARS-CoV-2? Am J Physiol Endocrinol Metab, doi:10.1152/ajpendo.00255.2020 (2020).

15 Lu Y1, Liu DX & JP., T. Lipid rafts are involved in SARS-CoV entry into Vero E6 cells. Biochem Biophys Res Commun 369, 6, doi: doi:10.1016/j.bbrc.2008.02.023 (2008).

16 Cao, X. et al. Spike protein of SARS-CoV-2 activates macrophages and contributes to induction of acute lung inflammation in male mice. FASEB J 35, e21801, doi:10.1096/fj.202002742RR (2021).

17 Jaishy, B. et al. Lipid-induced NOX2 activation inhibits autophagic flux by impairing lysosomal enzyme activity. J Lipid Res 56, 546–561, doi:10.1194/jlr.M055152 (2015).

18 Zang, H., Mathew, R. O. & Cui, T. The Dark Side of Nrf2 in the Heart. Front Physiol 11, 722, doi:10.3389/fphys.2020.00722 (2020).

19 Surviladze, Z., Sterk, R. T., DeHaro, S. A. & Ozbun, M. A. Cellular entry of human papillomavirus type 16 involves activation of the phosphatidylinositol 3-kinase/Akt/mTOR pathway and inhibition of autophagy. J Virol 87, 2508–2517, doi:10.1128/JVI.02319-12 (2013).

20 Feng, Z., Xu, K., Kovalev, N. & Nagy, P. D. Recruitment of Vps34 PI3K and enrichment of PI3P phosphoinositide in the viral replication compartment is crucial for replication of a positive-strand RNA virus. PLoS Pathog 15, e1007530, doi:10.1371/journal.ppat.1007530 (2019).

21 Jaber, N. et al. Class III PI3K Vps34 plays an essential role in autophagy and in heart and liver function. Proc Natl Acad Sci U S A 109, 2003–2008, doi:10.1073/pnas.1112848109 (2012).

22 Shen, B. et al. Proteomic and Metabolomic Characterization of COVID-19 Patient Sera. Cell 182, 59–72 e15, doi:10.1016/j.cell.2020.05.032 (2020).

23 Song, J. W. et al. Omics-Driven Systems Interrogation of Metabolic Dysregulation in COVID-19 Pathogenesis. Cell Metab 32, 188–202 e185, doi:10.1016/j.cmet.2020.06.016 (2020).

24 Schwarz, B. et al. Cutting Edge: Severe SARS-CoV-2 Infection in Humans Is Defined by a Shift in the Serum Lipidome, Resulting in Dysregulation of Eicosanoid Immune Mediators. J Immunol 206, 329–334, doi:10.4049/jimmunol.2001025 (2021).

25 Wu, D. et al. Plasma metabolomic and lipidomic alterations associated with COVID-19. Natl Sci Rev 7, 1157–1168, doi:10.1093/nsr/nwaa086 (2020).

26 Barberis, E. et al. Large-Scale Plasma Analysis Revealed New Mechanisms and Molecules Associated with the Host Response to SARS-CoV-2. Int J Mol Sci 21, doi:10.3390/ijms21228623 (2020).

27 Casari, I., Manfredi, M., Metharom, P. & Falasca, M. Dissecting lipid metabolism alterations in SARS-CoV-2. Prog Lipid Res 82, 101092, doi:10.1016/j.plipres.2021.101092 (2021).

28 Li, G. et al. Follow-up study on serum cholesterol profiles and potential sequelae in recovered COVID-19 patients. BMC Infect Dis 21, 299, doi:10.1186/s12879-021-05984-1 (2021).

29 Klann, K. et al. Growth Factor Receptor Signaling Inhibition Prevents SARS-CoV-2 Replication. Mol Cell 80, 164–174 e164, doi:10.1016/j.molcel.2020.08.006 (2020).

30 Yuen, C. K. et al. Suppression of SARS-CoV-2 infection in ex-vivo human lung tissues by targeting class III phosphoinositide 3-kinase. J Med Virol 93, 2076–2083, doi:10.1002/jmv.26583 (2021).

31 Nemazanyy, I. et al. Defects of Vps15 in skeletal muscles lead to autophagic vacuolar myopathy and lysosomal disease. EMBO Mol Med 5, 870–890, doi:10.1002/emmm.201202057 (2013).

32 Qin, Q. et al. Nrf2-Mediated Cardiac Maladaptive Remodeling and Dysfunction in a Setting of Autophagy Insufficiency. Hypertension 67, 107–117, doi:10.1161/HYPERTENSIONAHA.115.06062 (2016).

33 Zang, H. et al. Autophagy Inhibition Enables Nrf2 to Exaggerate the Progression of Diabetic Cardiomyopathy in Mice. Diabetes 69, 2720–2734, doi:10.2337/db19-1176 (2020).

34 Kobayashi, S. & Liang, Q. Autophagy and mitophagy in diabetic cardiomyopathy. Biochim Biophys Acta 1852, 252–261, doi:10.1016/j.bbadis.2014.05.020 (2015).

35 Ouyang, C., You, J. & Xie, Z. The interplay between autophagy and apoptosis in the diabetic heart. J Mol Cell Cardiol 71, 71–80, doi:10.1016/j.yjmcc.2013.10.014 (2014).

