## Supplemental Figures for "The Spike protein of SARS-CoV-2 impairs lipid metabolism and increases susceptibility to lipotoxicity: implication for a role of Nrf2"

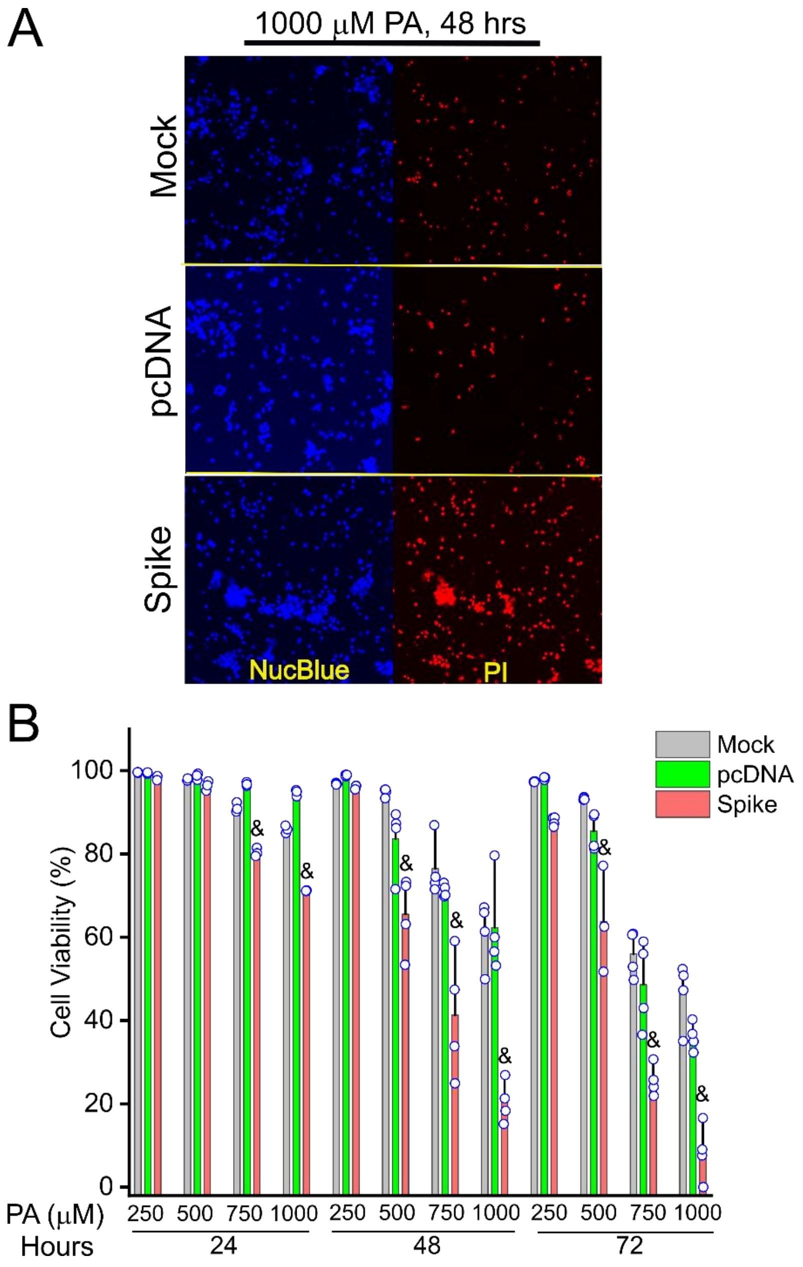


**Suppl Fig 1** PA-induced Spike protein-exaggerated lipotoxicity showed a dose- and time-dependent manner. &, indicates $p<0.05$ in the Spike cells as compared with the mock and pcDNA control cells for cell viability.

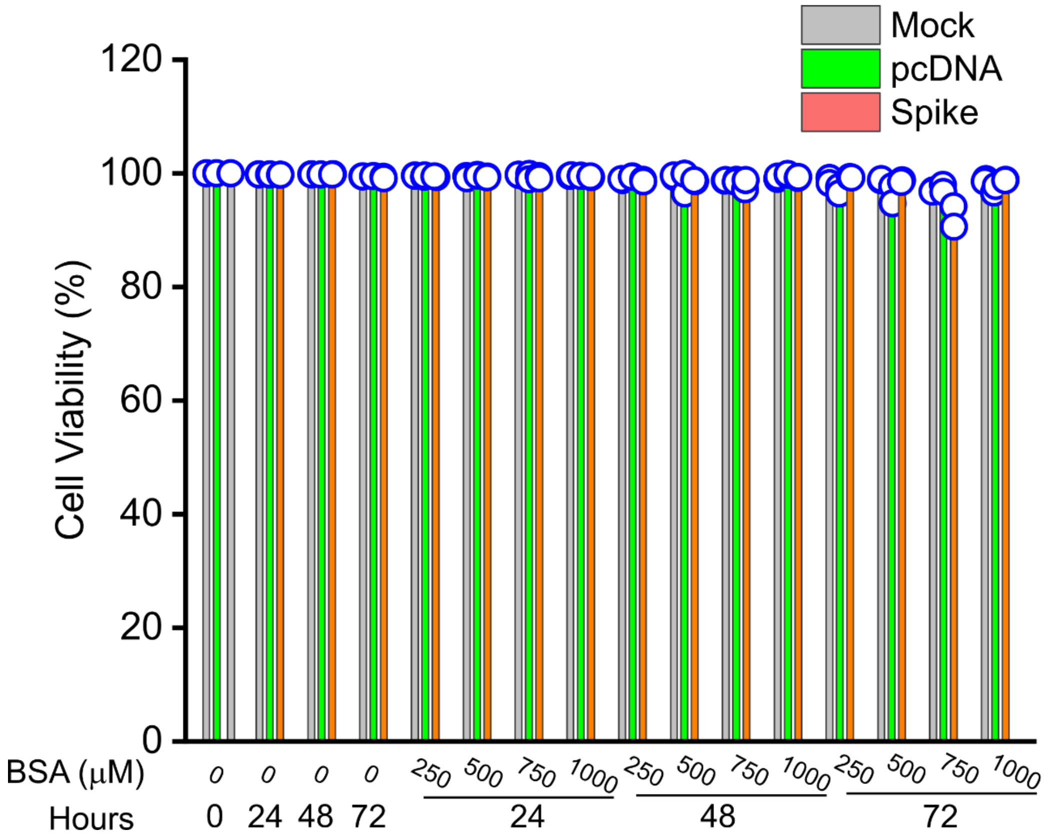


**Suppl Fig 2** The BSA with various concentrations doesn't cause significant cell death among the mock, pcDNA, and Spike cells.

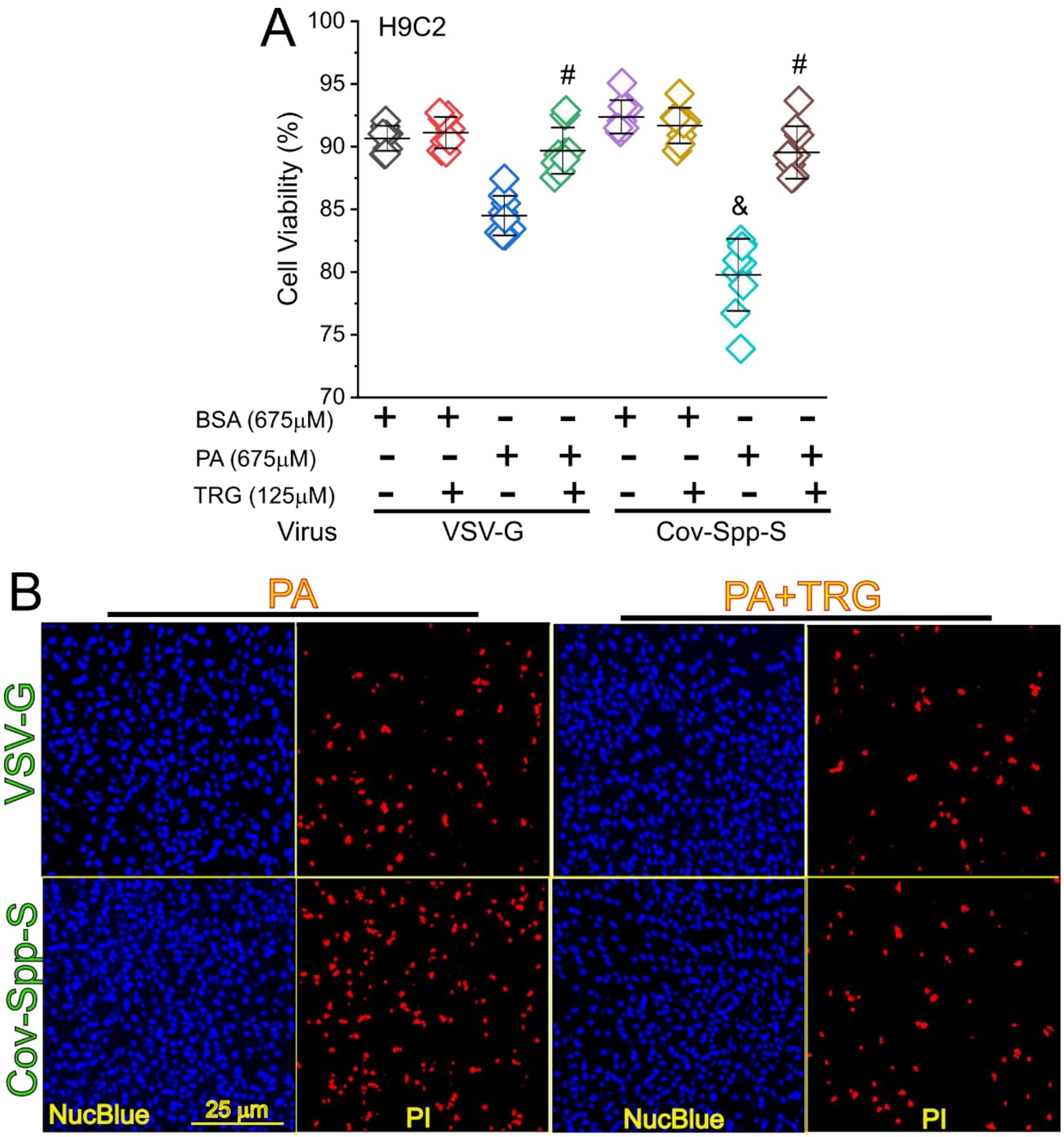


**Suppl Fig 3** TRG attenuated the Spike protein exaggerated PA-induced necrosis in H9C2 cells. (A) Quantitative data showing the cell viability of H9C2 cells after being infected by VSV-G or Cov-Spp-S lentivirus followed by PA treatment with or without TRG. BSA was used as a treatment control. &, indicates $p<0.05$ in the Cov-Spp-S-infected cells as compared with the VSV-G-infected cells upon PA treatment. #, indicates $p<0.05$ in the cells treated with PA plus TRG as compared with the cells treated with PA only. (B) Fluorescent images showing the cell viability in the VSV-G- or Cov-Spp-S-infected H9C2 cells after PA treatment with or without TRG in DMEM containing 5% FBS. The cells were co-stained by Propidium Iodide (PI) and Hoechst 33342 NucBlue Live Cell Stain dye to show the dead and live cells.
